# ANI, Mash and Dashing equally differentiate between *Klebsiella* species

**DOI:** 10.1101/2021.11.05.467470

**Authors:** Julie E. Hernández-Salmerón, Gabriel Moreno-Hagelsieb

## Abstract

Species of the genus *Klebsiella* are among the most important multidrug resistant human pathogens, though they have been isolated from a variety of environments. Given the need for quickly and accurately classifying newly sequenced *Klebsiella* genomes, we compared 982 *Klebsiella* genomes using different species-delimiting measures: Average Nucleotide Identity (ANI), which is becoming a standard for species delimitation, as well as Mash, Dashing, and DNA compositional signatures, which can be run in a fraction of the time required to run ANI. ROC analyses showed equal quality in species delimitation for ANI, Mash and Dashing (AUC: 0.99), followed by DNA signatures (AUC: 0.96). The groups obtained at optimal cutoffs were largely in agreement with species designation. Using optimized cutoffs, we obtained 17 species-level groups using either ANI, Mash, or Dashing, all containing the same genomes, unlike DNA signatures which broke the dataset into 38 groups. Further use of Mash to map species after adding draft genomes to the dataset also showed excellent results (AUC: 0.99), producing a total of 28 *Klebsiella* species in the publicly available genome collection. The ecological niches of *Klebsiella* strains were found to neither be related to species delimitation, nor to protein functional content, suggesting that a single *Klebsiella* species can have a wide repertoire of ecological functions.

## INTRODUCTION

Multidrug resistant bacteria represent a global threat to human health. Species of *Klebsiella* are among the most common antibiotic resistant human pathogens, causing as much as 50% mortality in infected neonatal, elderly and immunocompromised patients (Podschun R., 1998; Xu et al., 2017). *Klebsiella* species are considered ubiquitous in the environment (i.e., water, soil and plants), commonly found in the mucous surfaces of mammals, and as insect symbionts (Pinto-tomás et al., 2009; Martínez-Romero et al., 2015; Davis and Price, 2016). Despite their isolation source, imprecise detection methods affect the identification of potentially pathogenic strains (Podder et al., 2014; Rodríguez-Medina et al., 2019). Particularly, nearly identical *K. pneumoniae* strains have been found to be almost as virulent as strains of a clinical origin, despite disparate sources of isolation (Struve and Krogfelt, 2004; Huang et al., 2016; Dantur et al., 2018). This often leads to classification difficulties and taxonomic biases (Long et al., 2017) and highlights the need for faster and accurate techniques for the identification of *Klebsiella* isolates from environmental sources with the potential to infect humans.

To date, it is still complicated to develop techniques to identify *de novo* sequenced *Klebsiella* genomes from environmental or human strains at the species level (Garza-Ramos et al., 2015; Long et al., 2017). Some methods include the use of PCR-based probes with gene markers, such as the Multi Locus Sequence Typing (MLST) methods selected to identify *K. variicola* strains (Garza-Ramos et al., 2015; Barrios-Camacho et al., 2019); hierarchical clustering tools to classify *K. pneumoniae*, associated with antibiotic resistance to *β*-lactamase compounds (Berrazeg et al., 2013); and pan-genomic analysis to redefine subspecies of *K. pneumoniae* as new species of this genus (Caputo et al., 2015). These techniques aimed for the identification of a particular species, but the current amount of genomic data requires the extensive study of all available information at complete genomic level.

Currently, the most common measure to delimit species seems to be Average Nucleotide Identity (ANI), which has been used to revise the taxonomy of prokaryotes (Richter and Rosselló-Móra, 2009; Jain et al., 2018). However, to improve the speed for analyzing large amounts of sequence data, other methods have been developed. Mash, is based on the construction of MinHash sketches, consisting of somewhat random samples of small oligonucleotides (normally 21 bp long), transformed into hashes that can be efficiently computed and compared (Ondov et al., 2016). Dashing is another program using a computer transformation of oligonucleotides to improve speed and produce results similar to those of Mash (Baker and Langmead, 2019). Methods based on compositional analyses, focusing on nucleotides no more than 4 bp long, have also been used to group genomes (Richter and Rosselló-Móra, 2009; Moreno-Hagelsieb et al., 2013; Hernández-González et al., 2018).

The present study aims to test the accuracy and efficiency of different methods in grouping *Klebsiella* species based on program-specific thresholds, and to determine if the genetic content is associated with the environment of isolation.

## METHODS

### Genomic data

A total of 11,615 sequences of *Klebsiella* genomes were obtained from NCBI’s RefSeq database (Haft et al., 2018), by May 2021. Of these, only 982 were complete, or closed, genomes. This study focused mainly on the complete genomes dataset. The sources of isolation were compiled from the information available both in RefSeq and PATRIC (Davis et al., 2020).

To compare genomes, we calculated Average Nucleotide Identity with the program fastANI v1.32 (Jain et al., 2018), and distances with Mash v2.2.2 (Ondov et al., 2016), Dashing v0.5.7 (Baker and Langmead, 2019) and DNA Signatures (Campbell et al., 1999). We ran fastANI with a fragment length size of 1,020 bp to comply with the most usual ANI calculations. For Mash, we used the sketch function and the default k-mer size of 21 nt, with 5,000 sketches, rather than the default of 1000, since increasing the number of sketches should improve the accuracy of the obtained similarities. For Dashing, we tested three distance calculations: mash, full-mash and, its default, Jaccard. We selected a sketch size option of 2^14^ (-S 14). We also used a k-mer size of 21 to make the results more comparable to those of Mash. All the options used for each program are shown in Table 1. DNA signatures were calculated using an *ad hoc* program, written in perl. We obtained DNA signature distances for di, tri, and tetra nucleotides as reported previously by Moreno-Hagelsieb et al. (2013).

**Table 1.**
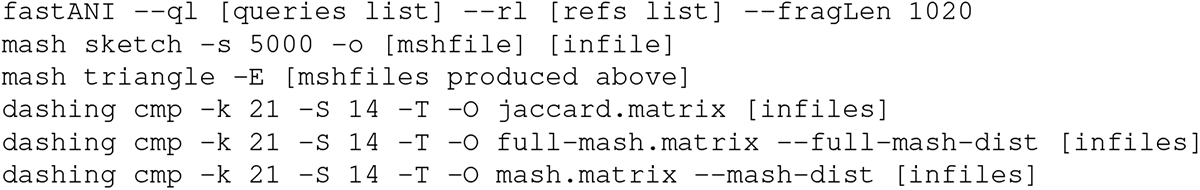
Commands used to run each program.

### Clustering analysis and optimal cut points

We performed Hierarchical Clustering to group *Klebsiella* species (Figure 1). The genomic distances required for clustering were calculated using the four programs: fastANI, mash, dashing, and DNA signatures as described above. ANI values were subtracted from 100 because they measure similarities, rather than distances. Distances between DNA signatures were Manhattan distances, divided by the length of the signature vector (Campbell et al., 1999; Moreno-Hagelsieb et al., 2013). Clusters were obtained using hclust and the divisive method (diana), implemented in R (R Core Team, 2021). Representative clusters were plotted using ggtree (Yu et al., 2017, 2018).

**Figure 1.**
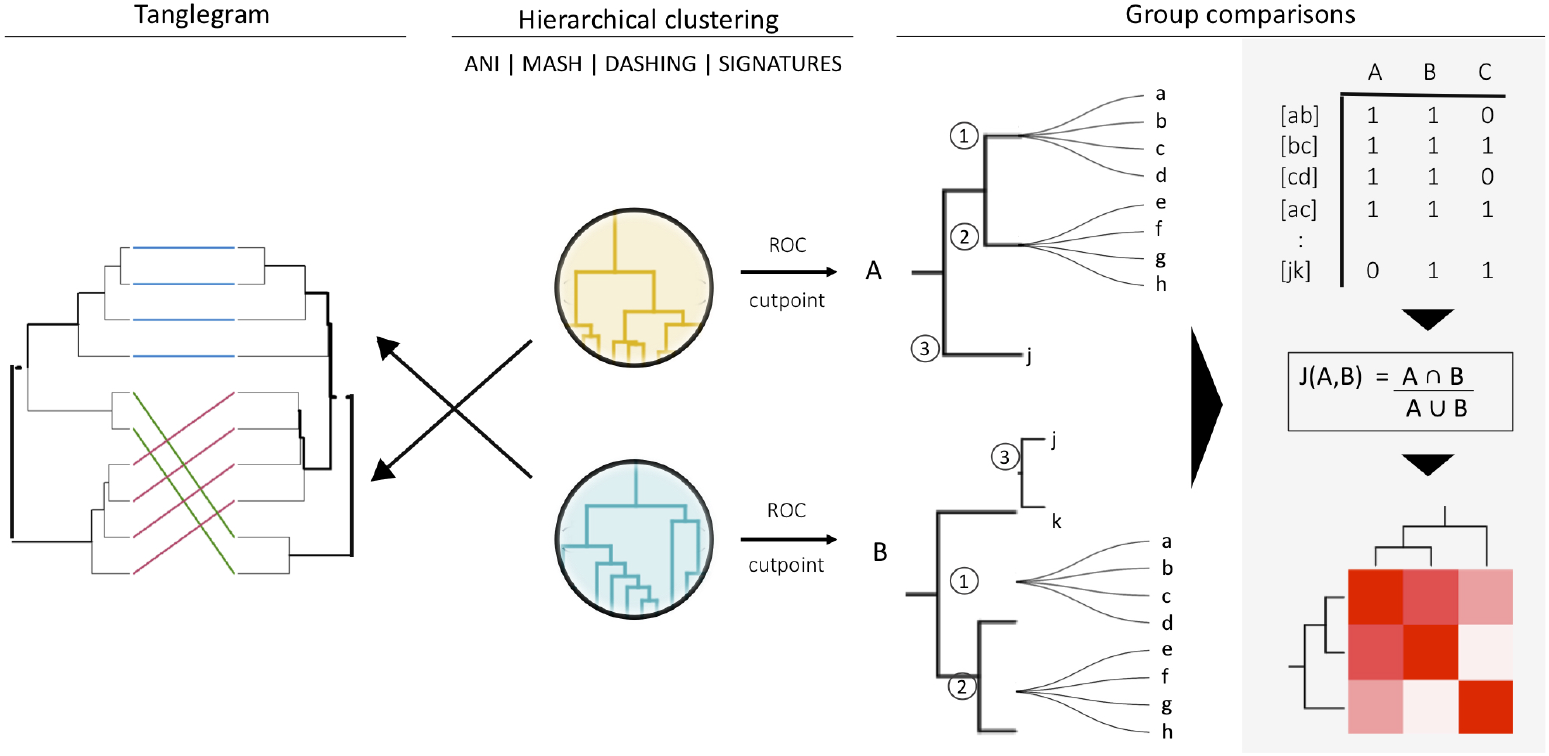
General strategy for programs and group comparisons. Hierarchical clustering was performed first (center). Clustering was followed by pairwise comparisons of all clusters obtained (tanglegram plots). Cut points specific for each dataset were calculated using a ROC curve analysis and the hierarchies were cut at that specific threshold. The members (a, b, c, etc.) of the resulting groups (nodes 1, 2 and 3) were rearranged into linked pairs and compared using Jaccard coefficients.

To evaluate and obtain optimum cutoff values for each of the programs tested, we calculated the ROC curves (Receiver Operating Characteristic). The positive dataset consisted of pairs of genomes assigned to the same species, while the negative dataset consisted of genomes in different species, but same genus. Both ROC analyses and optimal cut points were performed with the cutpointr R package (Thiele and Hirschfeld, 2021). We optimized for a maximum F1 score ([2 × *TP*]/[2 × *TP* + *FP* + *FN*]).

### Comparing groups

Hierarchical clusters were compared using a measure that evaluates the ‘entanglement’ between a pair of dendrograms, which is represented in a tanglegram plot (Figure 1). We used the dendextend package in R (Galili, 2015). The more similar the clusters are to each other, the lower the entanglement value, which ranges between 0 and 1.

To compare the groups resulting from cutting these hierarchical clusters at optimal cut points, we organized each group obtained into pairs of genomes belonging to the same group. The similarities between assigned pairs by each program were measured using Jaccard distances (Figure 1). An *ad hoc* program was written to analyse the species composition per group and compare them among the different programs. To avoid biased measures in the ROC and optimal cutpoints analyses due to an over-representation of the highly abundant species of *K. pneumoniae*, we performed a sub-sampling in which we selected five random samples containing 152 genomes of *K. pneumoniae*, each sample along with the rest of the complete genomes made up the five datasets.

### Domain content

Protein domains were obtained by comparing the proteins encoded by all genomes analysed, using mmseqs2 (Steinegger and Söding, 2017), against the appropriately formatted Pfam (Finn et al., 2015) and CDD databases (Marchler-Bauer et al., 2017). An *ad hoc* python3 script was written to gather the domain results into a single table to compare genome-to-genome domain content using Jaccard distances as implemented in the philentropy package in R (Drost, 2018). The results were used to perform divisive hierarchical clustering for an overall view of domain content similarities.

## RESULTS

### ANI, Mash and Dashing classify *Klebsiella* into almost identical species-level groups

A total of 982 complete genome sequences labeled as *Klebsiella* were downloaded from the RefSeq database (Haft et al., 2018). This dataset included 10 named species and 16 strains without a species designation, henceforth referred to as *Klebsiella* sp. (Figure 2).

**Figure 2.**
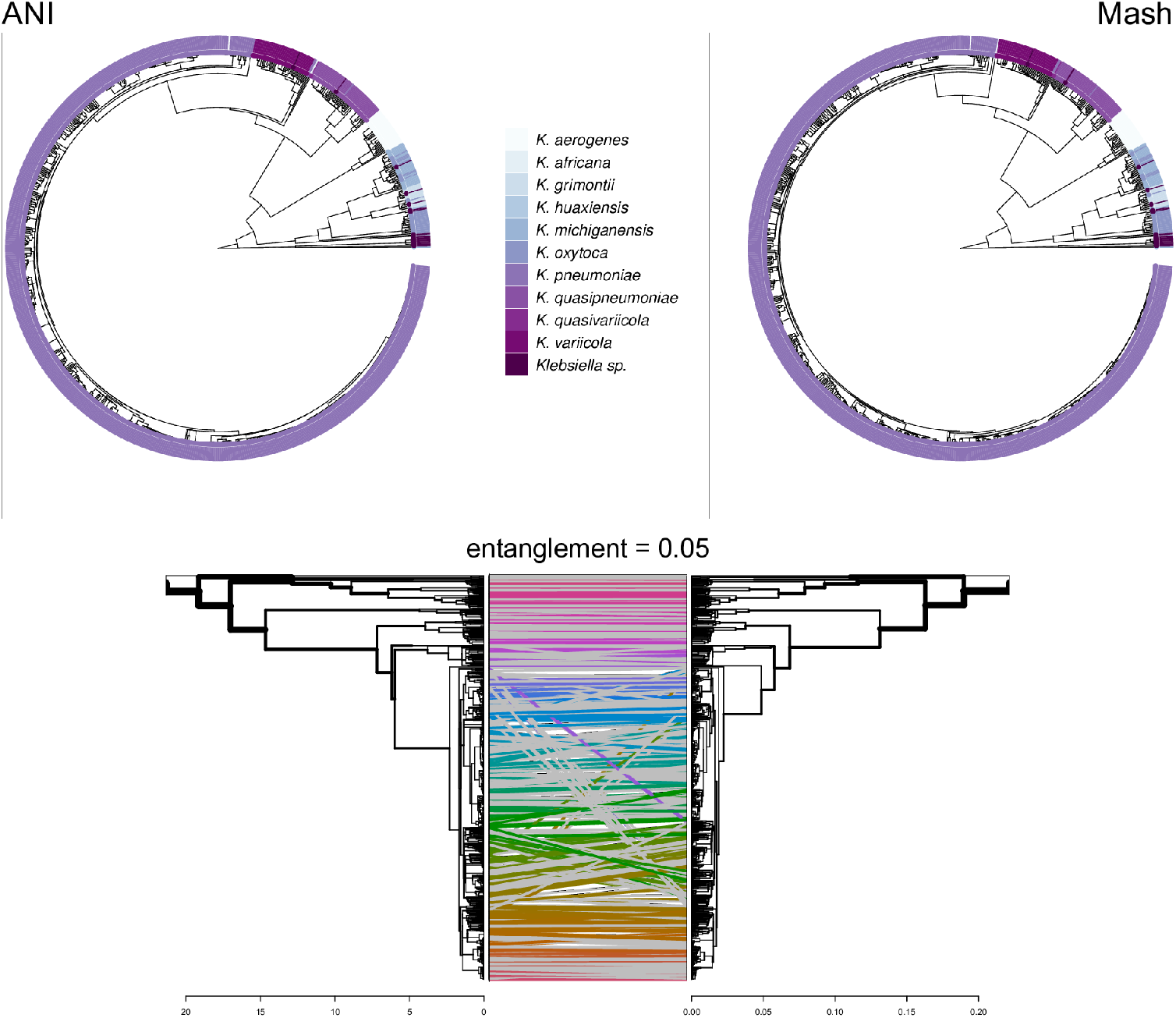
Top: Hierarchical clustering with distances calculated using ANI and Mash. Only assigned species as complete genomes were utilized. Bottom: Cluster comparison shows an entanglement value of 0.05, the closer the value to 0 the more similar the dendrograms are.

We used the distance/similarity calculated with Mash, DNA signatures and ANI, plus three available in Dashing (mash, full-mash and Jaccard), to produce hierarchical clusters for the complete genomes dataset. Figure 2 shows the representative clusters obtained with ANI and Mash; all the clusters are included in Supplementary F1. Entanglement values were calculated for all programs (Supplementary T1, F2), with ANI and Mash displaying the lowest entanglement score: 0.05 (Figure 2).

To test and compare the accuracy of the methods in species assignment, we performed Received Operating Characteristic (ROC) area under the curve (AUC) analyses. ANI, Mash and Dashing showed the same AUC of 0.996 (Figure 3), which means that these programs are close to perfection in distinguishing genomes from the same species from those in the same genus but different species. DNA signatures did not work as well, but had a respectable AUC of 0.956 (Figure 3).

**Figure 3.**
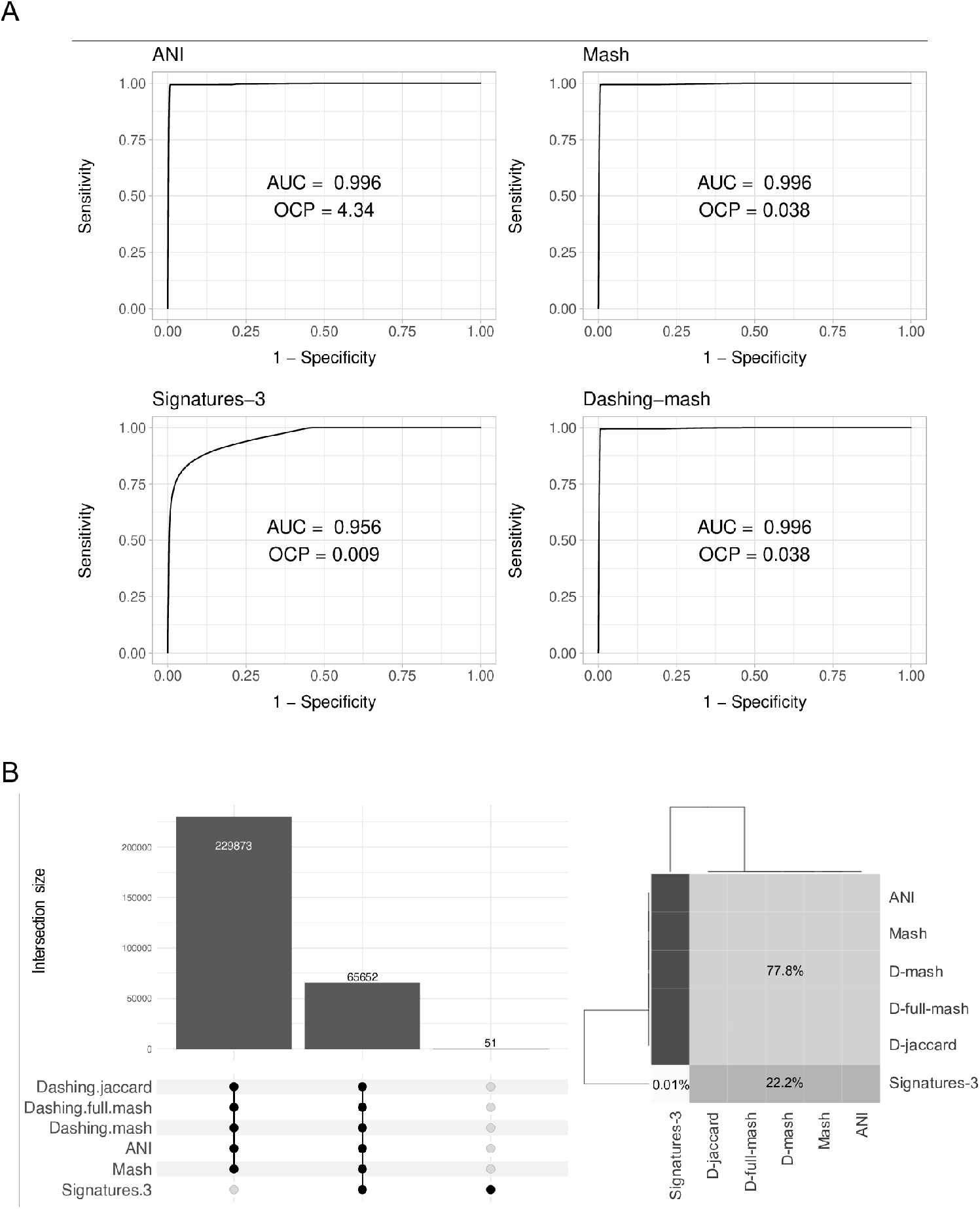
ROC curves analyses. **(A)** The area under the curve (AUC) suggests almost perfect delineation of species for ANI, Mash and Dashing mash, with a lower performance for DNA signatures. **(B)** The UpSet plot represents the genomes found in the same group after pruning the dendograms at the specific optimal cut point (OCP) obtained for each program. The heatmap shows that ANI, Mash and Dashing (full, mash-full and Jaccard) share 77% of the pairs found to belong to the same species, while they share only 22% with DNA signatures.

To compare results from cutting the clusters into species-level groups, we obtained cut points optimizing the F1 score. After pruning the trees using the specific cutoffs obtained, we found 17 groups of *Klebsiella* using the complete genomes dataset with ANI, Mash and Dashing, and 38 groups with DNA Signatures. To determine whether they contained the same groups, we checked each pair of genomes found within each group. Both the UpSet plot and the Jaccard distances (Figure 3B), showed that the Dashing algorithms, Mash and ANI shared 100% of the genome pairs obtained, while DNA signatures grouped 51 genome pairs in disagreement with the other programs.

The analysis of the composition of the 17 groups determined by ANI, Mash and Dashing showed the same distribution of the species *K. quasipneumoniae, K. quasivariicola, K. variicola, K. africana, K. grimontii* and *K. huaxiensis*, whose members were all grouped in single clusters using the program-specific cutoffs. Likewise, *K. pneumoniae, K. aerogenes* and *K. michiganensis* were equally split in three clusters, while the strains of *K. oxytoca* appeared in four groups, mostly sharing groups with members of *K. michiganensis* species (Supplementary F3). Further sub-sampling (n=5) of randomly selected datasets containing 152 genomes of *K. pneumoniae* plus all the genomes from the original dataset also showed the same results, an AUC of 0.99 for Mash, Dashing and ANI, while DNA signatures maintained a 0.95 of accuracy for all the five sub-samples. This time, the pruned clusters gave 14 species-level groups for ANI, Mash and Dashing; while DNA signatures produced different numbers of groups (35,38 and 39). Regardless the number of groups, ANI, Mash and Dashing showed equal composition of species per cluster in all the five datasets. However, a few differences were observed, Mash broke *K. aerogenes* in two clusters, unlike ANI and Mash who maintained the three clusters as the complete unique dataset. As for *K. pneumoniae*, whose members were also split in three groups before, when reducing the redundancy of genomes, they appeared split in only two groups similarly using ANI, Mash and Dashing (Supplementary F4).

### Neither species, nor protein content, seem related to isolation sources

To determine the relationship between genetic traits and the source of isolation in clustering species, we tested the four programs and clustered the complete genomes of *Klebsiella* according to the different hosts and the environment where they were isolated. The variety of hosts and source information available for the complete genomes was binned into five environments: animals, food, human, plants and water. Our results show that there is no clear evidence that the environment has a boundary effect to distinctively group *Klebsiella* strains (Figure 4).

**Figure 4.**
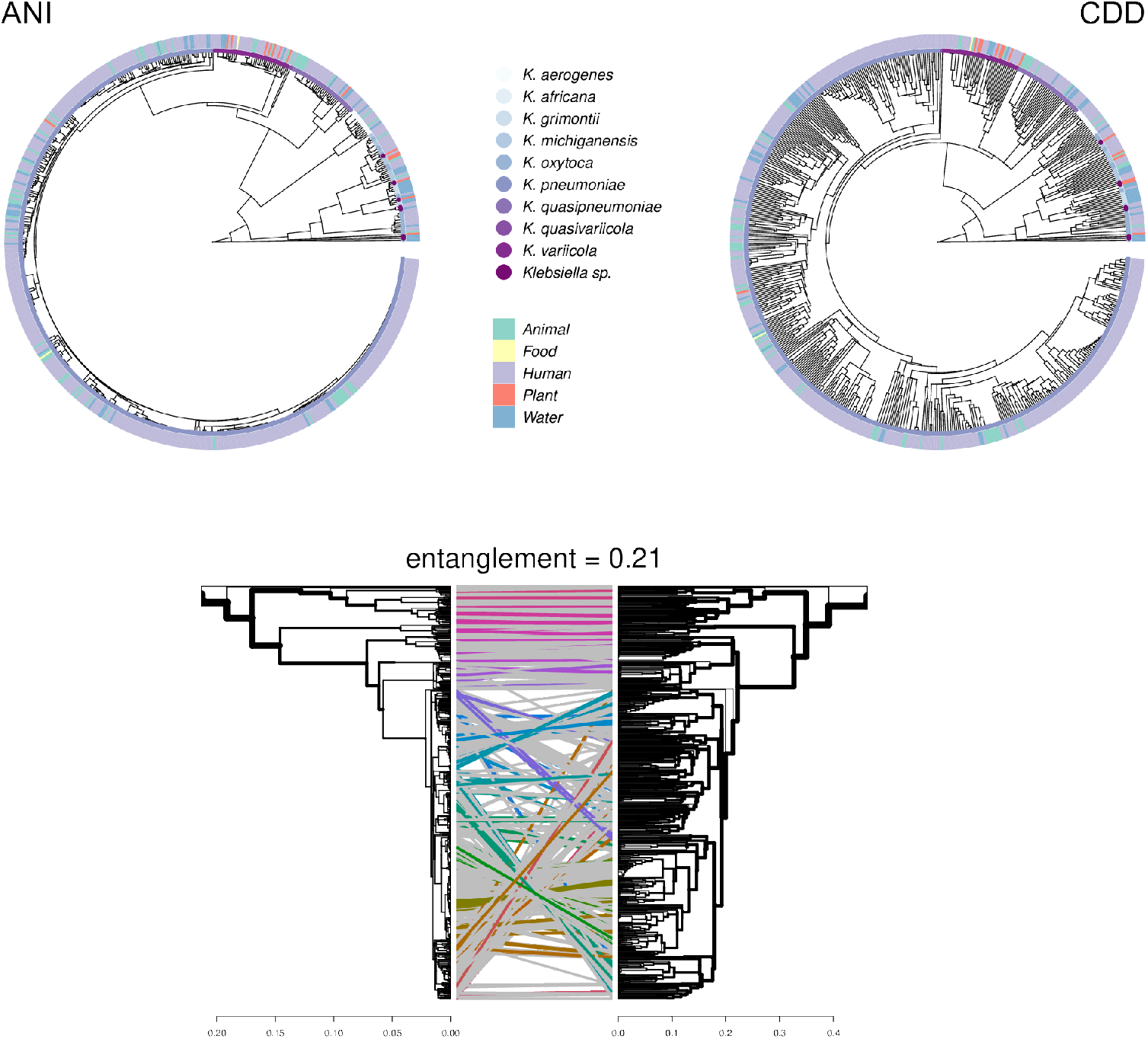
Hierarchical clustering using ANI and Jaccard distances between CDD protein domains annotations. Only genomes with isolation source annotations in PATRIC (Davis et al., 2020) are shown. The ANI and CDD clusters show a low entanglement value.

No matter where the strains were isolated from, they kept a clear species-specific cohesion, particularly those isolated from a clinical environment. In this regard, some authors have argued that intraspecific ecological and genetic interactions may constrain the diversification within a species (Cohan, 2019). Thus, environmental microbes isolated from different sources tend to display a genetic continuum, such as *K. variicola*, which was the most heterogeneous species found in almost all environments, yet it appears as a single cluster. Less cohesion is observed between *K. michiganensis* and *K. oxytoca*, which are divided into three and four groups, respectively. Thus, more specific thresholds and classification techniques might still be needed for classifying these strains. Overall, *K. aerogenes* appeared well clustered, except for *K. aerogenes* NCTC9644, which consistently grouped with *K. pneumoniae* strains, with, for example, ANI values are higher than 99%. Thereby we suggest to reconsider *K. aerogenes* NCTC9644 as a strain of *K. pneumoniae*.

### After adding draft genomes, Mash divides *Klebsiella* genomes into 28 species

In addition to the 982 complete *Klebsiella* genomes, we obtained a total of 10,633 draft genomes of different assembly status/categories: Chromosome, Scaffold and Contig (Supplementary T2). The length of the genomes comparing complete and draft genomes is shown in Figure 5, and from the different types of assembly in Supplementary F5. The genome lengths appeared similar regardless of the status of assembly. *K. oxytoca, K. michiganensis* and *K. grimontii* showed the largest genomes with over 7 Mbp, while strains of *K. aerogenes* had the smallest genomes ranging from 4.5 to 6 Mbp.

**Figure 5.**
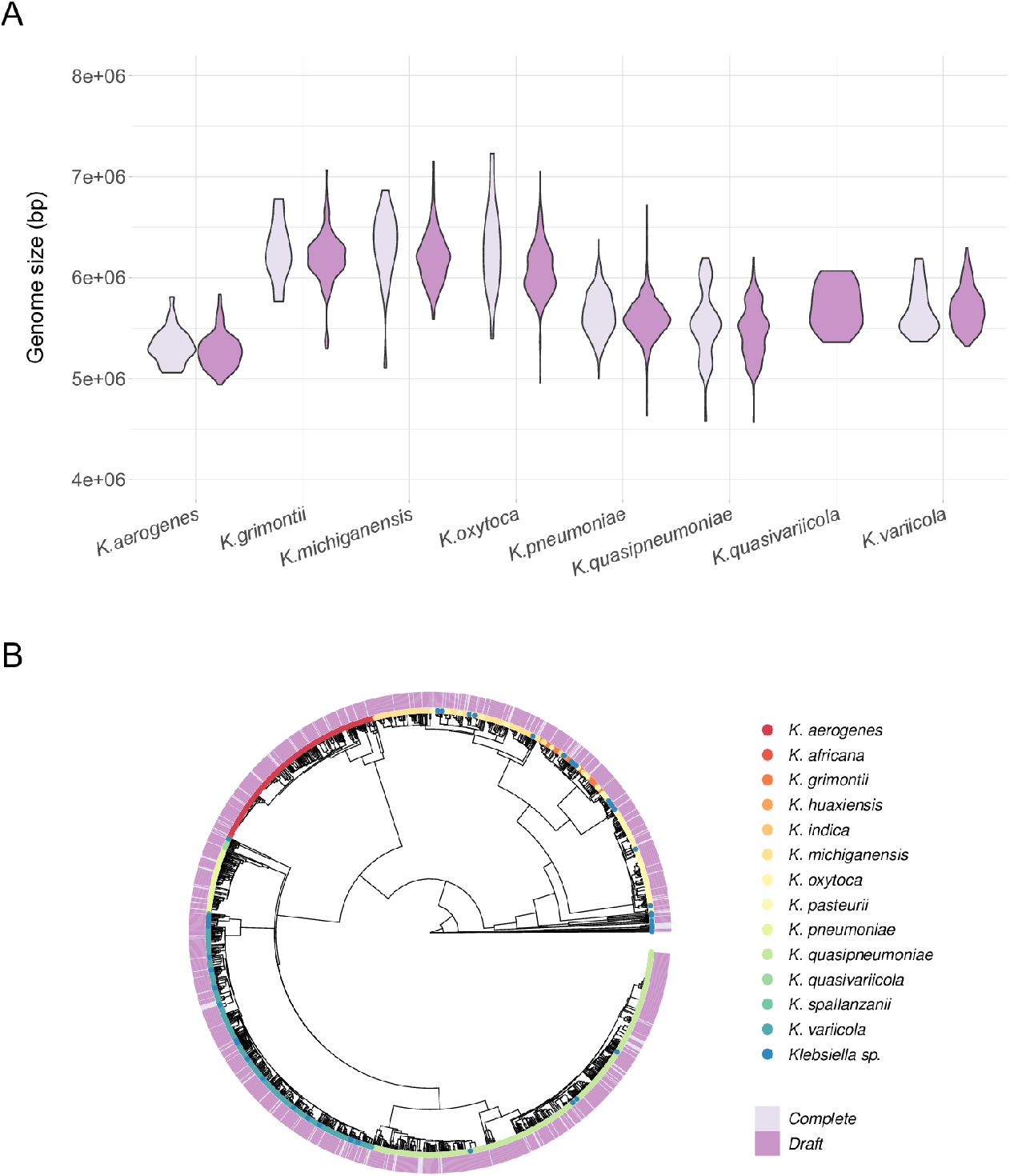
Draft genomes analysis.**(A)** Mash clustering for complete and draft genomes show highly similar results to those of obtained with the complete genomes alone. Representative tree using 1% of non-redundant *K. pneumoniae* genomes at the same cutoff of 0.038. **(B)** Genomic length of complete and draft genomes of *Klebsiella* species. The genome length appeared similar regardless the type of assembly. K. *oxytoca, K. michiganensis* and *K. grimontii* show the largest genomes with over 7 Mb, while *K. quasipneumoniae* has smaller genomes ranging from 4.5 to 6 Mbp.

Since Mash and Dashing produced the most similar results to ANI, we decided to test Mash with all types of genome assemblies and determine the reproducibility of the group assignments obtained with the complete genomes. We calculated a new cut point value from a new ROC analysis combining both datasets: complete and draft genomes, and obtained the same value of 0.038 as the previous for the complete genomes dataset. Using this cutoff, we obtained 28 species-level groups of *Klebsiella* (Figure 5). Our results showed that, regardless of the levels of assembly, Mash gives well separated groups with the same cutoff obtained using complete genomes.

## DISCUSSION

With the advances in sequencing technologies, it becomes essential to develop and validate efficient tools to handle large amounts of genomic data. ANI has been used worldwide since the authors found that a threshold of 95% mirrored the 70% of DNA-DNA hybridization technique with the advantage of being applicable across any sequenced prokaryotic species. (Richter and Rosselló-Móra, 2009). However, it is computationally intensive to calculate ANI, since genomic comparisons are based on pairwise alignments and calculations (Zhou et al., 2020). Apart from this, a recent revision has argued against the accuracy of the threshold proposed by the authors as a universal microbial species delineation, substantial sampling and species redundancy was found to alter the results and found no evidence of a universal genetic boundary among microbial species currently annotated in the NCBI taxonomy (Murray et al., 2021).

While recent developed programs have tried to overcome such inconsistencies, we are still not sure about the accuracy of the results they produce to definitely adopt new methods as standards and the best thresholds to assign species. Therefore, we need to evaluate the existing options that best resolve our specific genomic data. Our results show that Mash and Dashing produce the same results as ANI. We also provided specific thresholds for each program, which, in the case of Mash, 0.038, closely corresponds with a previous analysis in *Escherichia coli*, where the authors reported a cutoff of 0.037 (Abram et al., 2021). As for Dashing, we don’t have any knowledge so far of being tested in a specific bacterial species, apart from the experiments performed when the program was first released (Baker and Langmead, 2019). However, since we found the same species classification as ANI and Mash, we anticipate it may provide reliable results with highly improved computational efficiency. DNA signatures still offer a good approximation to the results produced by ANI and Mash, with an accuracy of 95.6%. However, little attention has been given to compositional methods, even though they have demonstrated various applications apart from clustering species (Richter and Rosselló-Móra, 2009), such as identification of exogenous DNA through horizontal transfer events, pathogenicity islands and bacteriophages (Bohlin, 2011).

Our study was conducted using all the genome sequences labeled as strains of *Klebsiella* in the RefSeq database. This genus contains a variety of species with diverse ecological functions. The most studied, *K. pneumoniae*, is considered the second leading pathogenic microorganism, after *E. coli. K. pneumoniae* are responsible for 70% of all nosocomial infections, along with other important opportunistic human pathogens, such as *K. aerogenes, K. variicola* and *K. oxytoca*. Members of the last two species also display plant-growth promoting activities such as nitrogen-fixation and production of relevant compounds with biotechnological applications (Tomulescu et al., 2021). The most recently described species, *K. huaxiensis, K. grimontii* and *K. africanensis* have been recovered from cattle and human feces/urine (Passet and Brisse, 2018; Hu et al., 2019; Rodrigues et al., 2019). The ubiquity of *Klebsiella* strains in natural environments, as well as their underestimated virulence potential, has posed a challenge to the proper identification and classification of *Klebsiella* species (Barrios-Camacho et al., 2019; Rodríguez-Medina et al., 2019). It has been estimated that between 2.5% and 10% of *K. pneumoniae* isolates are misidentified *K. variicola* strains (Rosenblueth et al., 2004; Fontana et al., 2019), which has led to fatal consequences for patients (Seki et al., 2013). This is an example of the significant implications that misidentification can have in epidemiology studies, which highlights the importance of a swift and adequate molecular identification that combines the appropriate computational tools and phenotypic approaches.

We extended our analysis to test data with less quality processed sequences since the majority of available genomes in public databases are unfinished. As of May 2021, only 7% (24,259/343,140) of the bacterial genomes in the RefSeq database are complete. We found that both sketching programs, Mash and Dashing, performed well for draft genomes without even altering the cutoff for the complete genomes analyzed. Mash has been previously reported to also perform well in whole metagenome comparisons (Dong et al., 2020) and to be very useful in fungal taxonomy (Gostinčar, 2020), along with many other promising applications (Zhou et al., 2020).

## CONCLUSIONS

Our results revealed that Mash and Dashing resolve species as accurately as ANI, but in a fraction of the time. We also provided specific thresholds for each program that performed well for this genus. The same threshold used for Mash worked for any genome assembly category. The isolation source was not found to be related to the species assignation, and the protein content did not show a strong correspondence with species or isolation source either. Therefore, only sequence similarities seem to define species boundaries so far. Further research on diverse bacterial species should be performed to have a broader perspective of the reliability and performance of these programs.

## ACKNOWLEDGMENTS

This work was supported by a Discovery Grant to GM-H from The Natural Sciences and Engineering Research Council of Canada (NSERC).

